# Algorithm optimization for weighted gene co-expression network analysis: accelerating the calculation of Topology Overlap Matrices with OpenMP and SQLite

**DOI:** 10.1101/2021.01.01.425026

**Authors:** Min Shuai, Xin Chen

**Affiliations:** School of Pharmacy, Chengdu University of T.C.M, Chengdu, China, 611137

## Abstract

**Motivation:** Weighted gene co-expression network analysis (WGCNA) is an R package that can search highly related gene modules. The most time-consuming step of the whole analysis is to calculate the Topological Overlap Matrix (TOM) from the Adjacency Matrix in a single thread. This study changes it to multithreading.

**Results:** This paper uses SQLite for multi-threaded data transfer between R and C++, uses OpenMP to enable multi-threading and calculates the TOM via an adjacency matrix on a Shared-memory MultiProcessor (SMP) system, where the calculation time decreases as the number of physical CPU cores increases.

**Availability and implementation:** The source code is available at https://github.com/do-somethings-haha/fast_calculate_TOM_of_WGCNA

## 1 Introduction

The weighted gene co-expression network analysis (WGCNA) package of R language can search for highly related gene modules (Langfelder and Horvath, 2008), and then find the potential genes in a certain metabolic pathway, or find the key genes in the gene regulatory network. It can be used for medical diagnosis (Wan *et al.*, 2018) or search for key genes related to a certain disease(Zhi *et al.*, 2018), and can study the mechanism of plant response to environmental stress (Chih-Ta *et al.*, 2019) or the structural genes and genes encoding transcription factors in a certain metabolic pathway (Lu *et al.*, 2019).

Changing the single-threaded algorithm to the multi-threaded algorithm is a major trend in algorithm development. In recent years, the increase in CPU single-core frequency has been limited, but the number of CPU cores has been increasing, and tools for parallel computing have gradually developed, such as OpenMP, CUDA, and MPI(Yang *et al.,* 2010). If single-threaded algorithms can be changed to multi-threaded algorithms, it will be extremely improve the calculation speed. For example, the single-threaded algorithm of sequence comparison has been changed to the multi-threaded algorithm (M *et al.,* 2017), and the algorithm of protein sequence search has been changed to the multi-threaded algorithm (Zhang *et al.,* 2016).

The <u>TOMsimilarity function</u> of the WGCNA package in R language that calculates the Topological Overlap Matrix (TOM) from the Adjacency Matrix is a single-threaded function, and the calculation takes too long, which hinders the use of this method in big data analysis.

This paper changes the algorithm of calculating TOM to a multi-threaded algorithm, which uses SQLite (https://www.sqlite.org/index.html) to realize multi-threaded high-speed data transmission between R and C++ languages, and uses OpenMP (https://www.openmp.org/) to realize multi-threaded calculation of TOM in C++ language. On a Shared-memory MultiProcessor (SMP) system, the calculation time decreases as the number of CPU cores used increases.

## 2 Methods

### 2.1 Data source

The test data used in this paper was downloaded from GSE61357 (EWipf *et al.,* 2014) of <u>Gene Expression Omnibus (GEO)</u>, which used gene chips to measure the expression of 30,677 genes. This paper only uses the expression matrix data of 19 samples whose sampling sites are root (GSM1502821~39).

### 2.2 Calculate Adjacency Matrix with WGCNA

All data of 19 samples from the test data are used in this paper, and the expression matrix is transposed so that the row names are sample names, the column names are the gene IDs.The soft threshold is set to 20. The Adjacency Matrix is calculated by the adjacency function of WGCNA package in R language.

### 2.3 Transfer Adjacency Matrix to C+ + from R

The 7G Adjacency Matrix is split into many data frames with a maximum column number of 2000, and these data frames are placed in a list. The foreach package in R language is used to start multiple processes, and each process obtains one data frame from the aforementioned list, while each process creates a connection to the SQLite database and creates a database file with one table. Every process writes a data frame to the corresponding table in the database file.

C++ language uses OpenMP to start multiple threads. Each thread connects to the corresponding database file, and reads the table in it. Finally, the Adjacency Matrix is stored in a two-dimensional array in C++ language.

### 2.4 Calculate TOM using C+ +

#### 2.4.1 convert the formulas

According to <u>the introduction of TOM in the WGCNA package</u>, the formulas for calculating TOM from the Adjacency Matrix are as follows.

##### 2.4.1.1 TOMType = “unsigned”, TOMDenom =“min”

If the parameters of TOMsimilarity function are set to TOMType = “unsigned”, and TOMDenom =“min”, then use the following formula

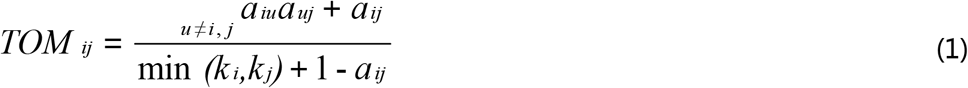

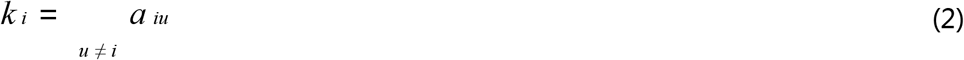

where *TOM* is the the Topological Overlap Matrix, *α* is the Adjacency Matrix, *i* is the row number of the Adjacency Matrix and TOM, *j* is the column number of the Adjacency Matrix and TOM. *u* increases from 1 to the maximum row number, also the maximum column number.

Because the Adjacency Matrix is a symmetric matrix, and the intermediate diagonals are all 1, we can deduce that

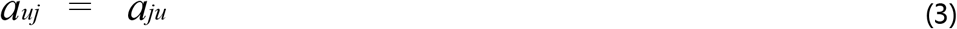

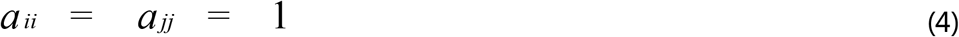

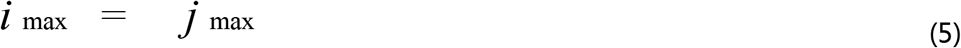

In combination with (2), (3) and (4), the formula (1) can be transformed into

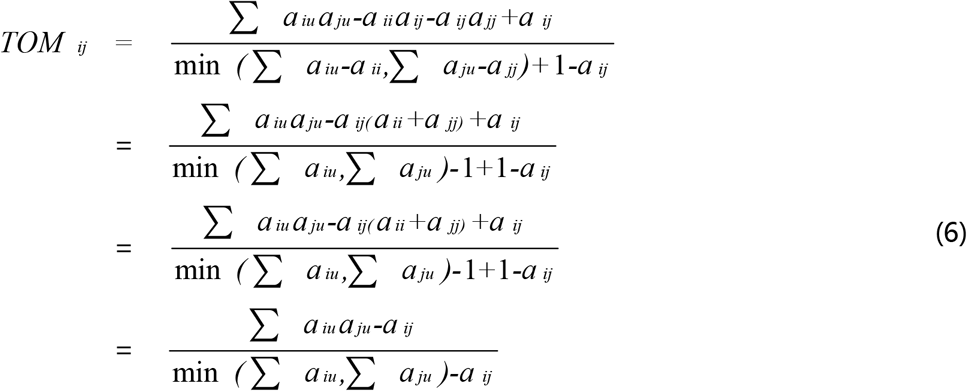

##### 2.4.1.2 TOMType = “unsigned”, TOMDenom = “mean”

If the parameters of TOMsimilarity function are set to TOMType =“unsigned”, and TOMDenom =“mean”, then use the following formula

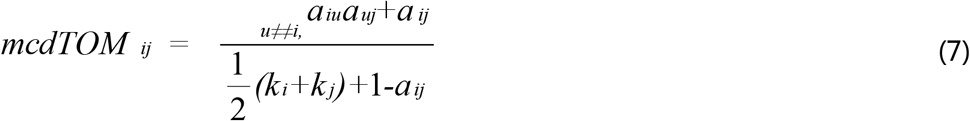

where *mcdTOM* is a more conservative modification of TOM.

In combination with (2), (3) and (4), the formula (7) can be transformed into

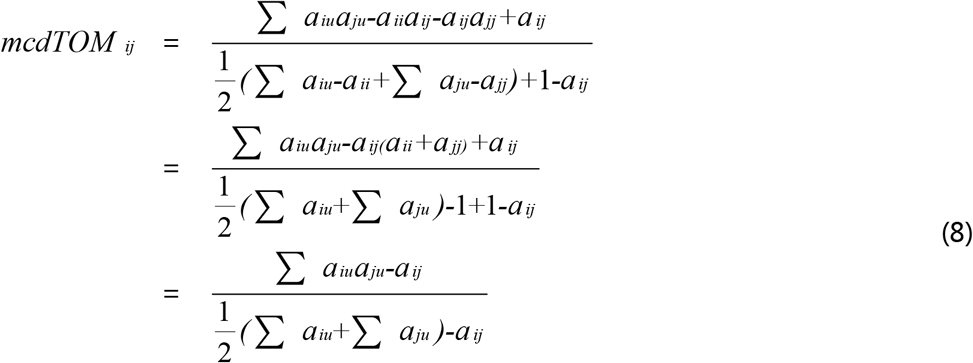

#### 2.4.2 Calculate the intermediate values

Calculating the *∑_a_iu_a_ju__* of the numerator in formula (6) or (8). According to the formula (6) or (8), the total number of intermediate values in the numerator is equal to the combination number: 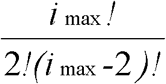. C++ language uses multiple threads to calculate these values and store them in a one-dimensional array.

Calculating *∑_a_iu__* and *∑_a_ju__* of the numerator in formula (6) or (8). C++ language uses multiple threads to calculate these values and then stores the values in a one-dimensional array.

#### 2.4.3 Calculate the final results

This paper uses the intermediate values to calculate each value of TOM according to (6) or (8).

### 2.5 Transfer TOM to R from C+ +

C++ language uses OpenMP to start multiple threads, and each thread connects to a database file. The TOM is split into multiple two-dimensional arrays with a maximum number of 2000. Every two-dimensional array is written into one table of the corresponding database file.

R language uses the foreach package to create multiple processes, and each process reads one of the aforementioned tables. Then, using the cbind function to bind the results of each process into a matrix named TOM.

## 3 Results

The results of the optimised multi-threaded algorithm are identical to the single-threaded results of the WGCNA package’s TOMsimilarity function, and they differ only in the last significant figures of the double-precision floating-point number due to the truncation error.

This paper tests the algorithms in shared memory multi-processor (SMP) system(CPU: Dual Intel Xeon Gold 5220 Processors, 36 cores in total. Memory: 8 × 32G, ECC, DDR4, 2666MHz, Operating system: Ubuntu 20.04). The WGCNA package’s TOMsimilarity function takes 180 minutes with a peak memory usage of about 10G. This paper’s algorithm takes 13 minutes if 72 threads are used with a peak memory usage of about 40G. The parallel acceleration ratio is 13.8, and the average acceleration ratio is 0.38. The relationship between the number of threads and the total time consumption is plotted in Figure 1.

**Fig. 1.**
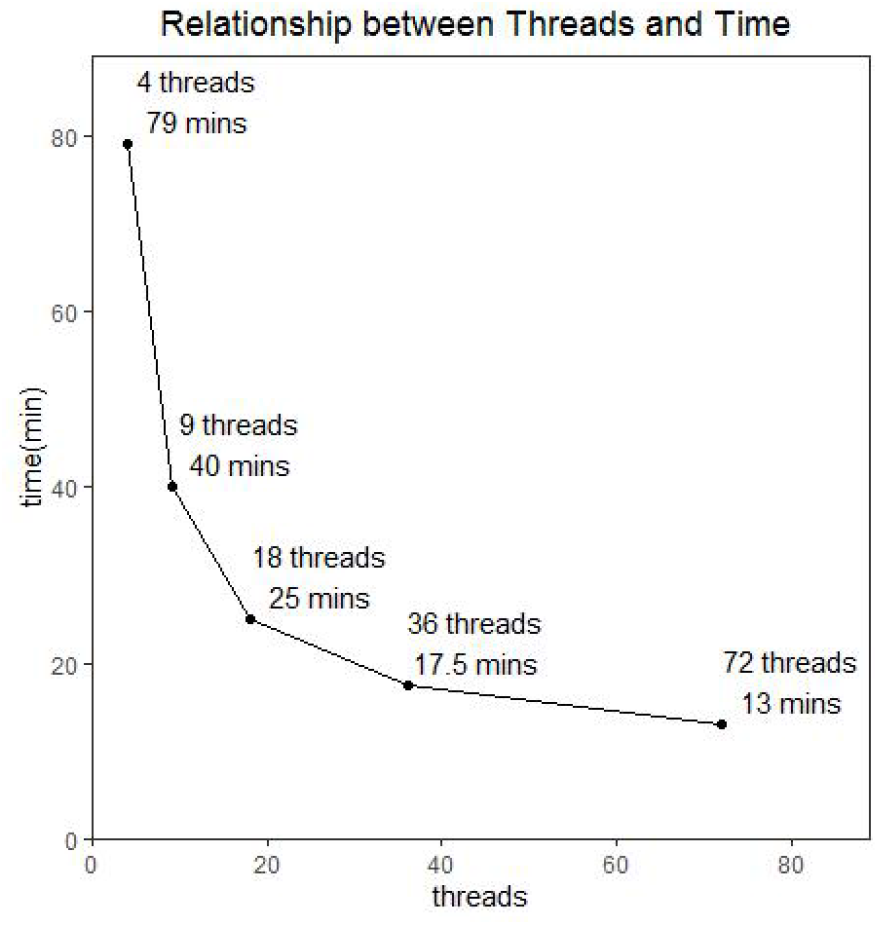
The relationship between the number of threads and the total time consumption for the optimized algorithm in this paper (CPU cores:36, total gene number: 30677). The WGCNA package’s single-threaded TOMsimilarity function takes 180 minutes.

## 4 Conclusions

There are a variety of clustering algorithms to search co-expressed genes, such as means of double clustering (Qinghui *et al.,* 2012), k-means, approximate kernel k-means, c-means, and hierarchical clustering (Uygun *et al.,* 2016). There are also many methods for constructing gene regulatory networks, however, the WGCNA package in R language is one of the best methods (Ruyssinck *et al.,* 2016). Since the release of the WGCNA package, there are lots of improvements, such as optimizing the degree of matching between gene modules and traits (David *et al.*, 2019).

If the algorithm of this paper is used to calculate the Adjacency Matrix with more than 10,000 genes, it is recommended to run on Linux systems, large-capacity ECC memory banks, server CPUs such as Intel Xeon series or AMD EPYC series, and good radiators.

The algorithm optimized in this paper greatly improves the calculation speed of the most time-consuming step in the WGCNA package. Thereby, the optimized algorithm can significantly improve the speed of the entire weighted gene co-expression network analysis, and it will enable the WGCNA package to be used more widely in the future, such as analyzing data from single cell sequencing (Luo *et al.*, 2015).

